# SCnorm: A quantile-regression based approach for robust normalization of single-cell RNA-seq data

**DOI:** 10.1101/090167

**Authors:** Rhonda Bacher, Li-Fang Chu, Ning Leng, Audrey P. Gasch, James A. Thomson, Ron M. Stewart, Michael Newton, Christina Kendziorski

**Author notes:** Corresponding author: Christina Kendziorski.

## Abstract

Normalization of RNA-sequencing data is essential for accurate downstream inference, but the assumptions upon which most methods are based do not hold in the single-cell setting. Consequently, applying existing normalization methods to single-cell RNA-seq data introduces artifacts that bias downstream analyses. To address this, we introduce SCnorm for accurate and efficient normalization of scRNA-seq data.

---

Protocols to quantify mRNA abundance introduce systematic sources of variation that obscure signals of interest; consequently, an essential first step in the majority of mRNA expression analyses is normalization, whereby systematic variations are adjusted for in an effort to make expression counts comparable across genes and/or samples. Within-sample normalization methods adjust for gene-specific features such as GC-content and gene length to facilitate comparisons across genes within an individual sample, whereas between-sample normalization methods adjust for sample-specific features such as sequencing depth to allow for comparisons of a gene’s expression across samples^1^. In this work, we present a method for between-sample normalization, although we note that the R implementation, R/SCnorm, also allows for adjustment of gene-specific features (**Supplementary Section S1**).

A number of methods are available for between-sample normalization in bulk RNA-seq experiments^2,3^. Although the details differ slightly among approaches, each attempts to identify genes that are relatively stable across cells, then uses those genes to calculate global scale factors (one for each sample applied commonly across genes in the sample) to adjust each gene’s read counts in each sample for sequencing depth. Although these methods demonstrate excellent performance for bulk RNA-seq, the abundance of zeros and increased technical variability present in scRNA-seq data compromise their performance in the single-cell setting^4^. Recent methods have been developed specifically for scRNA-seq normalization that accommodate both an abundance of zeros and increased technical variability^5,6^, however, like many bulk methods, they too calculate global scale factors.

Although existing methods for scRNA-seq normalization show considerable improvement over bulk approaches, they are unable to accommodate a major bias that to date has been unobserved in scRNA-seq data. Specifically, scRNA-seq data show systematic variation in the relationship between read counts and sequencing depth (referred to hereinafter as the count-depth relationship) that is not accommodated by a single scale factor common to all genes in a cell. For highly expressed genes, counts increase directly with increases in depth, similar to most genes in a bulk RNA-seq experiment. That is not the case for moderate and lowly expressed genes, where counts typically increase at a slower rate than expected with increases in depth (**Fig. 1 and Supplementary Figure S1).** Global scale factors adjust for a count-depth relationship that is assumed common across genes. When this is not the case, normalization via global scale factors leads to over-correction for lowly and moderately expressed genes and, in some cases, under-normalization of highly expressed genes (**Fig. 1**). As a result, normalization methods that rely on global scale factors are not appropriate for scRNA-seq.

**Figure 1:**
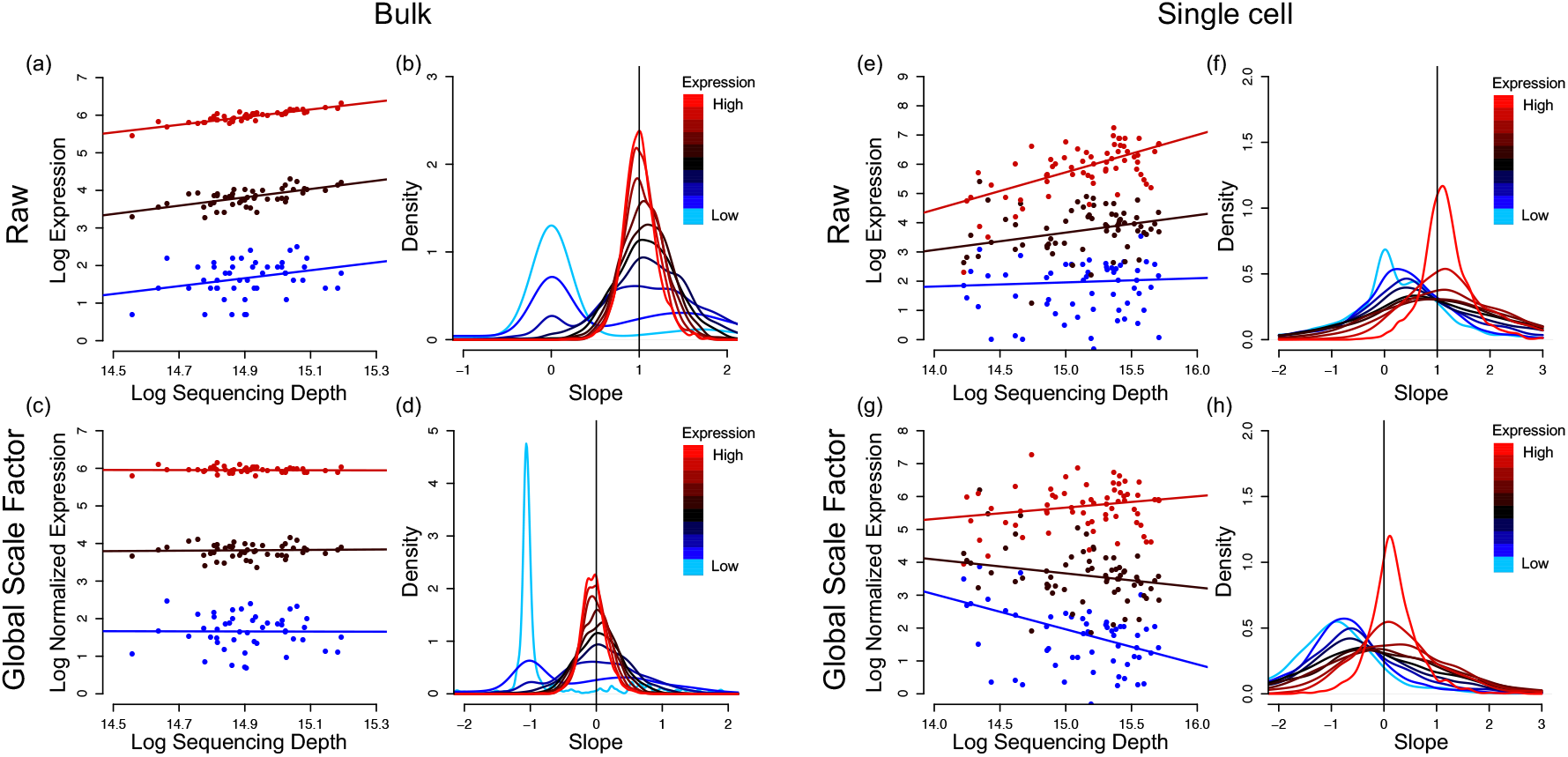
For each gene, median quantile regression was used to estimate the count-depth relationship before normalization and after normalization via MR for the H1 bulk RNA-seq data set (panels **(a) – (d)**) and the DEC scRNA-seq data set (panels **(e)-(h)**). Panel **(a)** shows log-expression vs. log-depth and estimated regression fits for three genes having low, moderate, and high expression defined as median expression among non-zero un-normalized measurements in the 10^th^-20^th^ quantile, 40^th^-50^th^ quantile, and 80^th^-90^th^ quantile, respectively. Panel **(b)** shows densities of slopes within each of ten equally sized gene groups where a gene’s group membership is determined by its median expression among non-zero un-normalized measurements. Panels **(c)** and **(d)** show the same data as panels **(a)** and **(b),** respectively, but here the data are normalized via MR. Panels **(e)-(h)** are structurally identical to **(a)-(d)** for the DEC scRNA-seq data set. Qualitatively similar results are observed if slopes are calculated via generalized linear models (**Supplementary Section S3** and **Supplementary Figure S1**).

To address this, we propose SCnorm for robust normalization of scRNA-seq data. Briefly, SCnorm uses quantile regression to estimate the dependence of read counts on sequencing depth for every gene. Genes with similar dependence are then grouped, and a second quantile regression is used to estimate scale factors within each group. Within-group adjustment for sequencing depth is then performed using the estimated scale factors to provide normalized estimates of expression. Although SCnorm does not require spike-ins, performance may be improved if good spike-ins are available **(Supplementary Section S2**).

SCnorm was evaluated and compared with MR^3^, transcripts-per-million (TPM)^7^, scran^5^, SCDE^8^, and BASiCS^6^. Because BASiCS requires spike-ins, results are only shown for data sets where spike-ins are available. In the simulation study, we assessed the ability with which normalized estimates of expression could be used to estimate fold-change as well as the sensitivity and specificity for identifying differentially expressed (DE) genes. The simulations vary with respect to assumptions and extent of DE, which should help to ensure a reasonably realistic evaluation of the operating characteristics of SCnorm.

In SIM I, two scenarios were considered where the number of groups of genes having different count-depth relationships (*K*) is set to 1 and 4, respectively. When *K* = 1, the relationship between expression and sequencing depth is similar across all genes, as in bulk RNA-seq. Each simulated data set contains two conditions, the second condition having approximately four times as many reads; 20% of the genes are defined to be DE where overall the DE is balanced (10% up-regulated and 10% down-regulated on average). Prior to normalization, counts in the second condition will appear 4 times higher on average given the increased sequencing depth. However, if normalization for depth is effective, fold-change estimates should be near one, and only simulated DE genes should appear DE. **Supplementary Figure S2a** shows that when *K =* 1, with the exception of TPM, fold-change estimates are consistently robust among methods, and all normalization methods provide data that results in high sensitivity and specificity for identifying DE genes (**Supplementary Fig. S2b**). However, when K = 4, only SCnorm maintains good operating characteristics, whereas global scale factor based approaches overestimate fold-changes for low to moderately expressed genes due to overcorrection of sequencing depth (**Supplementary Fig. S2c, d**).

In SIM II, counts are generated as in Lun *et al.* 2016^5^, following their simulation study scenarios 1, 2, 3, and 4. Briefly, scenario 1 contains no DE genes; scenarios 2, 3, and 4 contain moderate DE, strong DE, and varying magnitudes of DE, respectively. **Supplementary Figure 3** shows that SCnorm is similar to *scran* with respect to fold change estimation; it also retains relatively high sensitivity and specificity for identifying DE genes (**Supplementary Fig. S4**).

To further evaluate SCnorm, we conducted an experiment that, similar to the simulations, sequenced cells at very different depths. Specifically, we used the Fluidigm C1 system to capture 92 H1 human embryonic stem cells (hESCs). Each cell’s fragmented, indexed cDNA was then split into two groups prior to pooling for sequencing. In the first group (referred to as H1-1M), indexed cDNA from cells was pooled at 96 cells per lane and in the second (H1-4M) cDNA from cells was pooled at 24 cells per lane, resulting in approximately 1 million and 4 million mapped reads per cell in the two groups, respectively. Prior to normalization, counts in the second group will appear four times higher on average given the increased sequencing depth. However, if normalization for depth is effective, fold-change estimates should be near one, and all genes should appear to be EE since the cells between the two groups are identical. **Figure 2 (a)** shows estimates of fold-changes calculated between the two groups. As shown, SCnorm provides normalized data that results in fold-change estimates near one, whereas other methods show biased estimates.

**Figure 2:**
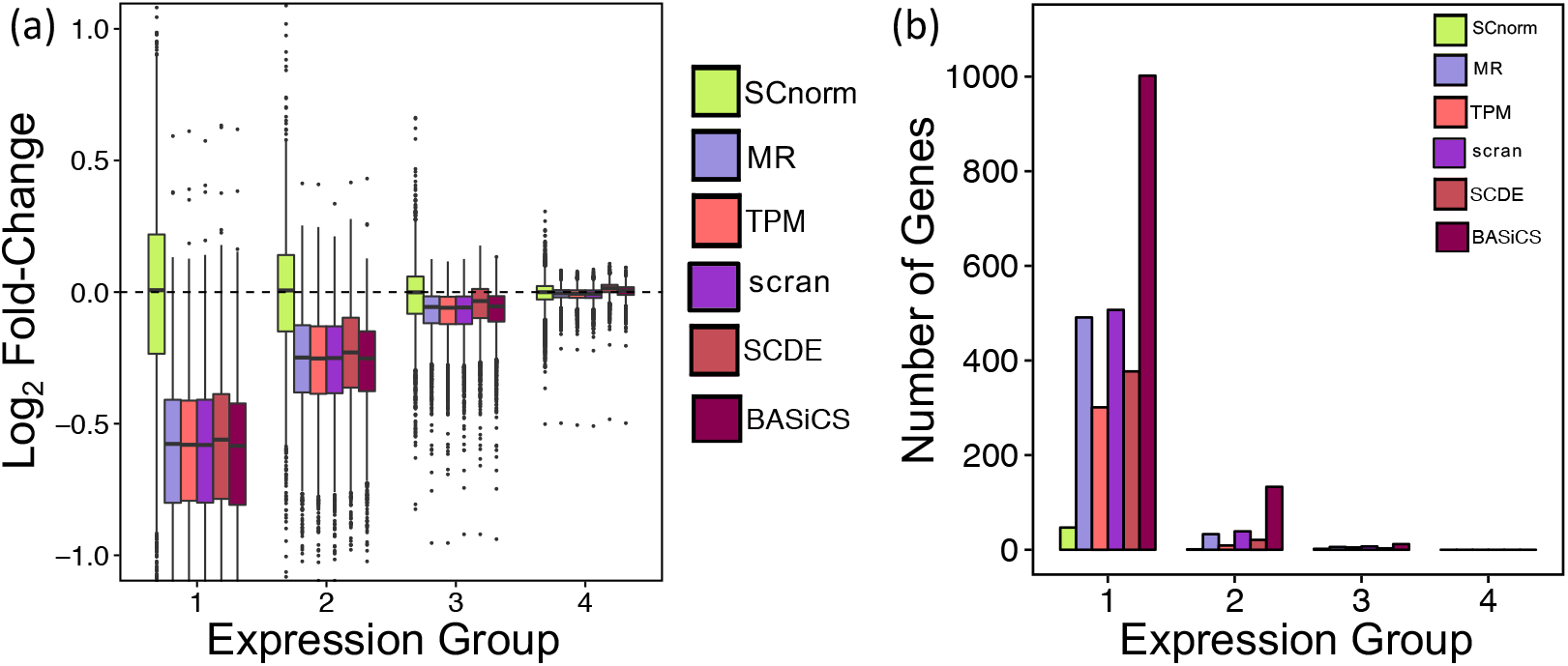
For each gene, the fold-change of non-zero counts between the H1-4M and H1-1M groups was computed for data following normalization via SCnorm, MR, TPM, scran, SCDE, and BASiCS. Box-plots of gene-specific fold-changes are shown in panel **(a)** for data normalized by each method. The number of genes identified as DE using MAST is shown in panel **(b)**. Genes are divided into four equally sized expression groups based on their median among non-zero un-normalized expression measurements and results are shown as a function of expression group. Motivation for considering non-zero counts to calculate fold-change is discussed in **Supplementary Section S4**.

To evaluate the extent to which biases introduced during normalization affect the identification of DE genes, we applied MAST (FDR = 0.05) to identify DE genes between the H1-1M and H1-4M conditions. Normalization with SCnorm resulted in the identification of 50 genes, whereas MR, TPM, scran, SCDE, and BASiCS resulted in 530, 315, 553, 401, and 1147 DE genes, respectively, being identified. The majority of DE calls made using data normalized from these latter approaches are lowly expressed genes (**Fig. 2 (b)**), which appear to be over-normalized (**Fig. 2 (a)**). Similar results were obtained when the experiment was repeated using H9 cells (**Supplementary Fig. S5**).

The performance of SCnorm was also evaluated on a number of case study data sets. For these evaluations, a data set was considered well normalized if the relationship between counts and depth was removed following normalization for all genes. **Figure 3** and **Supplementary Figures S6-S12** demonstrate that SCnorm provides for robust normalization of scRNA-seq data when the count-depth relationship is common across genes, as in a bulk RNA-seq experiment (or a deeply sequenced scRNA-seq experiment); and that SCnorm outperforms other approaches when this relationship varies systematically, as in a typical scRNA-seq experiment.

**Figure 3:**
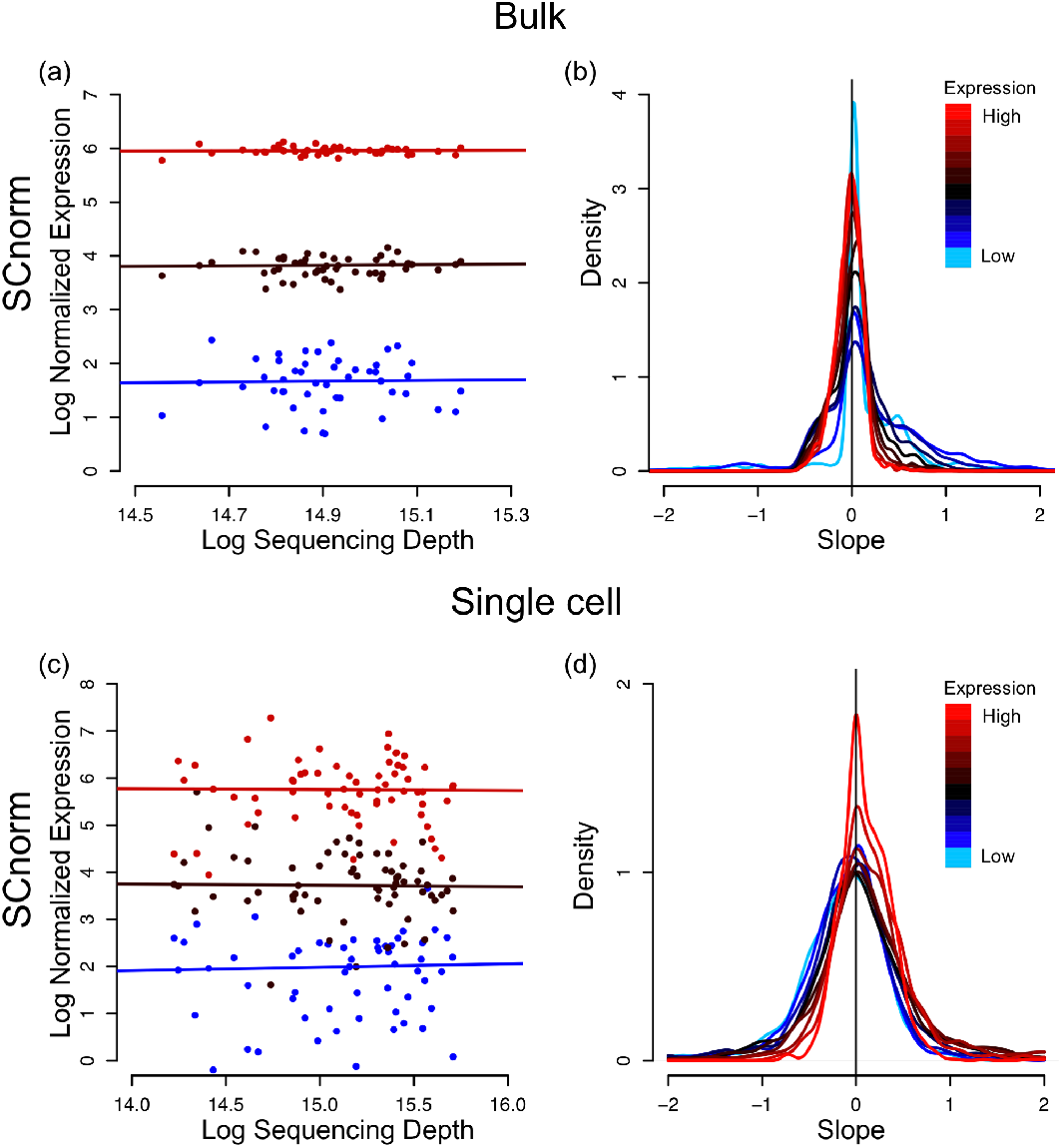
For each gene, median quantile regression was used to estimate the count-depth relationship after normalization via SCnorm for the H1 bulk RNA-seq data set (panels **(a) – (b)**) and the DEC scRNA-seq data set (panels **(c)-(d)**). Results are structurally identical to those shown in Figure 1, but with data normalized by SCnorm. Qualitatively similar results are observed if slopes are calculated via generalized linear models (**Supplementary Section S3** and **Supplementary Figure S6**).

The scRNA-seq technology offers unprecedented opportunity to address biological questions, but accurate data normalization is required to ensure meaningful results. Our approach allows investigators to accurately normalize data for sequencing depth, and consequently to improve downstream inference.

## ACCESSION CODES

The H1 bulk and the H1-1M, H1-4M, H9-1M, H9-4M case study datasets will be released at the gene expression omnibus (GSE85917) upon acceptance of the manuscript.

## ACKNOWLEDGMENTS

This work was supported by NIH GM102756, NIH U54 AI117924, and the Morgridge Institute for Research. We thank J. Bolin, A. Elwell, and B.K. Nguyen for the preparation and sequencing of the RNA-seq samples and P. Jiang and S. Swanson for performing the RNA-seq read processing.

## AUTHOR CONTRIBUTIONS

R.B. and C.K. designed the research, developed the method, and wrote the first version of the manuscript. L.C. performed experiments and quality control on scRNA-seq data generated from H1 and H9 hESCs. R.B. analyzed all datasets. L.C., N.L., A.P.G., R.S. and M.N. analyzed results from early versions of the method which helped during method refinement. All authors contributed to writing the manuscript.

## ONLINE METHODS

### Filter

Genes without at least 10 cells having non-zero expression were removed prior to all analyses. They are not shown in plots.

### SCnorm

SCnorm requires estimates of expression, but is not specific to one approach. Estimates may be obtained via RSEM^7^, HTSeq^9^, or any method providing counts per feature. Let *Y*_*g,j*_ denote the log non-zero expression count for gene *g* in cell *j* for *g* = 1,…, *m* and *j* = 1,…, *n*; *X*_*j*_ denote log sequencing depth for cell *j.* Motivation for considering non-zero counts is provided in **Supplementary Section S4**.

The number of groups for which the count-depth relationship varies substantially, *K*, is chosen sequentially. SCnorm begins with *K* = 1. For each gene, the gene-specific relationship between log expression and log sequencing depth is represented by 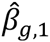 using median quantile regression with a first degree polynomial: *Q*^0.5^(*Y*_*g,j*_|*X*_*j*_) = *β*_*g*,0_ + *β*_*g*,1_*X*_*j*_. The overall relationship between log expression and log sequencing depth for all genes in the *K* = 1 group is also estimated via quantile regression. Since the median might not best represent the full set of genes within the group, and since multiple genes allow for estimation of somewhat subtle effects, in this step SCnorm considers multiple quantiles t and multiple degrees *d*:

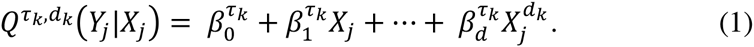

The specific values of *τ*_*k*_ and *d*_*k*_, 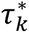 and 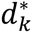, are those that minimize 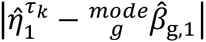, where 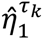 represents the count-depth relationship among the predicted expression values as estimated by median quantile regression using a first degree polynomial: 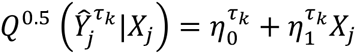. Scale factors for each cell are defined as 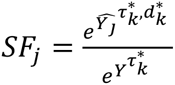 where 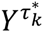 is the *τ*^\**th*^ quantile of expression counts in the *k*^th^ group. Normalized counts 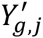 are given by 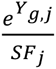.

To determine if *K* = 1 is sufficient, the gene-specific relationship between log normalized expression and log sequencing depth is represented by the slope of a median quantile regression using a first degree polynomial as detailed above. *K*= 1 is considered sufficient if the modes of the slopes within each of 10 equally sized gene groups (where a gene’s group membership is determined by its median expression among non-zero un-normalized measurements) are all less than 0.1. Any mode exceeding 0.1 is taken as evidence that the normalization provided with *K* = 1 is not sufficient to adjust for the count-depth relationship for all genes and, consequently, *K* is increased by one and the count-depth relationship is estimated within each of the *K* groups using equation (1). For each increase, the *K*-medoids algorithm is used to cluster genes into groups based on 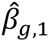; if a cluster has fewer than 100 genes, it is joined with the nearest cluster.

When multiple biological conditions are present, SCnorm is applied within each condition and the normalized counts are then re-scaled across conditions. During rescaling, all genes are split into quartiles based on median expression among non-zero un-normalized measurements. Within each group and condition, each gene is scaled by a common scale factor defined as the median of the gene specific fold-changes between each gene’s condition-specific mean and the gene-specific mean across conditions, where means are calculated over non-zero counts. Motivation for considering non-zero counts during re-scaling is discussed in **Supplementary Section S4**.

### SCnorm.SI

SCnorm does not require spike-ins, since we find that the performance of spike-ins in scRNA-seq is often compromised (**Supplementary Fig. S13-S14**), and many labs do not use them for normalization^10^. However, if good spike-ins are available, performance of SCnorm may be improved in the post-normalization scaling step, which is required when multiple conditions are available. Recall that in SCnorm, during rescaling, all genes are split into quartiles based on median expression among non-zero un-normalized measurements. In SCnorm.SI, the same is done with spike-ins and, if the spike-ins are representative of the full range of expression, we expect them to be approximately evenly divided among the four groups. Within each group and condition, each gene is scaled by a common scale factor defined as the median of the spike-in specific fold-changes between each spike-in’s condition-specific mean and the spike-in’s specific mean across conditions, where means are calculated over non-zero counts. For more on SCnorm.SI, see **Supplementary Section S2**.

### Application of comparable methods

All analyses were carried out using R version 3.3.0 unless otherwise noted. The method MR, originally described by Anders and Huber^3^, was implemented using the DESeq R package version 1.24.0 using the default settings of the *estimateSizeFactorsForMatrix* function. TPM estimates were obtained as output from RSEM version 1.2.3. The method scran was implemented with the scran R package version 1.0.0; size factors were obtained using the function *computeSumFactors*. The pool sizes were set to 5, 10, 15, and 20; and size factors were constrained to be positive. SCDE was implemented in R version 3.2.2 using the SCDE R package version 1.99.1 with default parameter settings, and normalized counts were obtained using the function *scde.expression.magnitude*. Finally, BASiCS was implemented using the BASiCS R package version 0.4.1 using R vesion 3.2.2, obtained from Github at <u>https://github.com/catavallejos/BASiCS</u>; and normalized expression estimates were obtained using the function *BASiCS_DenoisedCounts* where *BASiCS_MCMC* was run with N = 20,000, Burn = 10,000, and default parameters used otherwise.

### Evaluation of methods

Gene-specific count depth relationships were estimated using median quantile regression as well as regression with a negative binomial generalized linear model (glm). The *quantreg* package in R was used with the Frisch–Newton interior point method to carry out the median regressions; MASS in R was used to fit the glms. Zeros are not included in the fits since our goal is to estimate the count-depth relationship present in data before and after normalization, and that relationship is obscured by dropouts, which are largely technical. Because glm's are sensitive to outliers, an initial glm to estimate the count-depth relationship is fit on the un-normalized data and the top two and bottom two residual gene expression values were removed from each gene prior to estimating the final count-depth relationship via glm. Since the same set of putative outliers were removed for every method, excluding these values will not bias results in favor of any one method.

MAST was used to identify DE genes, using the MAST R package version 0.933, obtained from Github at <u>https://github.com/RGLab/MAST</u>. The continuous component test was considered and differential zeros were not used to evaluate performance of normalization methods since all normalization methods leave zeros un-normalized. P-values from MAST were adjusted using Benjamini & Hochberg^11^. Unless otherwise noted, a DE gene was defined as one with corrected p-value < 0.05, which controls the false discovery rate at 5%. ROC curves were plot using the R package *ROCR*. The false positive and true positive rates were calculated by *ROCR*, with a positive representing a DE gene. Average ROC curves show the average true positive rate.

### Simulation SIM I

Data were simulated to match characteristics of the H1-1M and H1-4M datasets. For each gene *g*, gene-specific intercepts 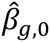, slopes 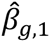, and variance intercepts 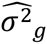 were estimated using median quantile regression on the H1-1M data. Two SIM I simulation scenarios were generated: *K* = 1 and *K* = 4. In the *K* = 1 simulations, only genes having at least 75% non-zero expression values and 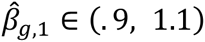 were used. For the *K* = 4 simulations, genes were split into four equally sized groups based on 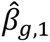. The medians of 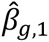 were calculated within each group; denote these by *β*_*med*,1_, *β*_*med*,2_, *β*_*med*,3_, and *β*_*med*,4_, respectively. For genes in the *k*^th^ group, genes having 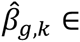 (*β*_*med,k*_ − 0.1, *β*_*med,k*_ + 0.1) were used, where *β*_*med,k*_ is the median 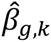 over all genes.

For a given gene, counts were simulated on the log scale as 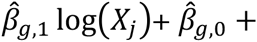 *ε*_*g,j*_ and then exponentiated, where 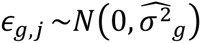. Two biological conditions were simulated: one condition with 90 cells simulated from sequencing depths ranging from 500,000 to 1.5 million reads (*X*_*j*_ was sampled uniformly between 500,000 and 1.5 million) and a second condition with 90 cells simulated with depths ranging from 2 to 6 million reads (*X*_*j*_ was sampled uniformly between 2 and 6 million). For a randomly selected set of cells, counts were set to zero, where the proportion set to zero was defined to match the proportion observed empirically. Each simulated dataset contained 1200 genes, 80% EE and 20% DE. For approximately half of the DE genes, fold-changes were sampled uniformly between 2 and 4, and counts in the second condition were multiplied by the sampled fold-change. The other (approximately) half of DE genes were simulated similarly, but with counts in the first condition multiplied by the sampled fold change to keep the DE balanced. **Supplementary Figure S15** shows that basic summary statistics are well preserved between the simulated and case study data.

### Simulation SIM II

Counts are generated as in Lun *et al.* 2016^5^ following their simulation study scenarios 1, 2, 3, and 4. In that simulation set up, three populations were simulated. We here consider populations 1 and 2.

### H1 bulk data

The dataset contains 48 samples of H1 hESCs as described in detail in Hou *et al.* 2015^12^. The H1 bulk RNA-seq data have an average sequencing depth of 3 million mapped reads per sample.

### H1 and H9 case studies

Undifferentiated H1 or H9 hESCs were cultured in E8 medium^13^ on Matrigel-coated tissue culture plates with daily media feeding at 37 °C with 5% (vol/vol) CO_2_. Cells were split every 3-4 days with 0.5 mM EDTA in 1 X PBS for standard maintenance. Immediately before preparing single cell suspensions for each experiment, hESCs were individualized by Accutase (Life Technologies), washed once with E8 medium, and resuspended at densities of 5.0-8.0 X 10^5^ cells/mL in E8 medium for cell capture. The H1 hESCs are registered in the NIH Human Embryonic Stem Cell Registry with the Approval Number: NIHhESC-10-0043. Details of the H1 cells can be found online (http://grants.nih.gov/stem_cells/registry/current.htm?id=29). The H9 hESCs are registered in the NIH Human Embryonic Stem Cell Registry with the Approval Number: NIHhESC-10-0062. Details of the H9 cells can be found online (http://grants.nih.gov/stem_cells/registry/current.htm?id=414). All the cell cultures performed in our laboratory have been routinely tested and have been found negative for mycoplasma contamination and authenticated by cytogenetic tests.

Single-cell loading, capture, and library preparations were performed following the Fluidigm user manual “*Using the C1 Single-Cell Auto Prep System to Generate mRNA from Single Cells and Libraries for Sequencing.*” Briefly, 5,000-8,000 cells were loaded onto a medium size (10-17 *μm*) C1 Single-Cell Auto Prep IFC (Fluidigm), and cell-loading script was performed according to the manufacturer’s instructions. The capture efficiency was inspected using EVOS FL Auto Cell Imaging system (Life Technologies) to perform an automated area scanning of the 96 capture sites on the IFC. Empty capture sites or sites having more than one cell captured were first noted and those samples were later excluded from further library processing for RNA-seq. Immediately after capture and imaging, reverse transcription and cDNA amplification were performed in the C1 system using the SMARTer PCR cDNA Synthesis kit (Clontech) and the Advantage 2 PCR kit (Clontech) according to the instructions in the Fluidigm user manual. Full-length, single-cell cDNA libraries were harvested the next day from the C1 chip and diluted to a range of 0.1-0.3 ng/µL. Diluted single-cell cDNA libraries were fragmented and amplified using the Nextera XT DNA Sample Preparation Kit and the Nextera XT DNA Sample Preparation Index Kit (Illumina). Libraries were multiplexed either at 24 or 96 single cell cDNA libraries per lane to target 4 or 1 million mapped reads per cell, respectively, and single-end reads of 67-bp were sequenced on an Illumina HiSeq 2500 system. We refer to the data obtained from 24 libraries per lane as the H1-4M set, since approximately 4 million mapped reads per cell were generated. For similar reasons, H1-1M is used to refer to the data obtained from 96 libraries per lane.

Reads were mapped against the Hg19 Refseq reference via Bowtie 0.12.8^14^ allowing up to two mismatches and up to 20 multiple hits. The expected counts and TPM’s were estimated via RSEM 1.2.3^7^. Cells having less than 5,000 genes with expected counts > 1 or those that upon inspection of cell images displayed doublets or appeared dead were removed in quality control. 92 H1 cells passed the quality control. 91 H9 cells passed quality control.

### Buettner case study^15^

Single cell RNA-seq expression data were downloaded from ArrayExpress E-MTAB-2805. In this experiment, *Mus musculus* embryonic stem cells were sorted using fluorescence-activated cell sorting (FACS) to determine cell cycle stage; cells were then captured using the C1 Fluidigm system. Libraries were multiplexed and sequenced across four lanes using an Illumina HiSeq 2000 system. Gene-level read counts were generated by HTSeq version 0.6.1. Here we consider the three data sets each having 96 cells in either G1, S, or G2M stage of the cell cycle. The data have average sequencing depths of 4.9, 6.5, and 4.5 million, respectively. Cells having sequencing depths less than 10,000 were removed prior to analysis which resulted in 95 G1, 88 S, and 96 G2M cells.

### Islam case study^16^

Single cell RNA-seq expression data were downloaded from GEO GSE29087. In this experiment, *Mus musculus* R1 embryonic stem cells (ES) and embryonic fibroblasts were captured using a semi-automated cell picker on a 96-well capture plate; libraries were generated using the STRT protocol and sequenced using on a Genome Analyzer IIx system. Gene-level counts were obtained by counting reads mapped using Bowtie^14^ for each feature. Here we consider two datsets, one having 48 ES cells and the other having 44 EF cells. The datasets have average sequencing depths of 180,000 reads and 800,000 reads, respectively.

### DEC case study

The dataset contains 64 H1 cells consisting of the first batch of experiments studying H1 differentiation towards definitive endodermal cells as described in detail in Chu *et al.* 2016^17^. The DEC scRNA-seq data have an average sequencing depth of 4 million mapped reads per cell. The data can be downloaded from GEO GSE75748.

